# The reserve of joint torque determines movement coordination

**DOI:** 10.1101/2021.04.12.439505

**Authors:** Germain Faity, Denis Mottet, Simon Pla, Jérôme Froger

## Abstract

Humans coordinate biomechanical degrees of freedom to perform tasks at minimum cost. When reaching a target from a seated position, the trunk-arm-forearm coordination moves the hand to the well-defined spatial goal, while typically minimising hand jerk and trunk motion. However, due to fatigue or stroke, people visibly move the trunk more, and it is unclear what cost can account for this. Here we show that people recruit their trunk when the torque at the shoulder is too close to the maximum. We asked 26 healthy participants to reach a target while seated and we found that the trunk contribution to hand displacement increases from 11% to 27% when an additional load is handled. By flexing and rotating the trunk, participants spontaneously increase the reserve of anti-gravitational torque at the shoulder from 25% to 40% of maximal voluntary torque. Our findings provide hints on how to include the reserve of torque in the cost function of optimal control models of human coordination in healthy fatigued persons or in stroke victims.

## 1. Introduction

When reaching to grasp an object from a seated position, the body segments that bring the hand closer to the target are the trunk, arm and forearm. This kinematic chain is redundant so that, theoretically, reaching the final posture can be achieved in an endless number of different patterns of coordination of the trunk, arm and forearm ^1^. However, empirically, the reaching movement quasi exclusively relies on shoulder flexion and elbow extension ^2^.

According to the optimal control framework, the CNS achieves goals at a minimal cost ^3^. To coordinate body segments, the CNS minimises both error and effort ^4^, where the error is task related and the effort includes internal costs ^5,6^, such as minimum energy expenditure, minimum muscle fatigue or minimum sense of effort ^7^. When reaching to grasp an object from a seated position, the CNS takes into account that the trunk is massive, with greater inertia than the upper arm and forearm, and with a larger lever arm that requires more torque to start and stop. Thus, when it can do otherwise, the CNS has no interest in recruiting the costly trunk within the coordination.

However, the CNS sometimes recruits the trunk. For example, the trunk is recruited when the target is too far away to be reached with the shoulder-elbow coordination ^8^. Obviously, to minimise the error, one has to pay the price of a higher effort. More surprisingly, the trunk is also recruited when the target is not far away, typically in people suffering from hemiparesis following a stroke ^9,10^, in children with and without cerebral palsy ^11^ or in fatigued healthy people ^12^. Peeters *et al*. (2018) argue that it may be easier to accurately perform a reaching task in children or patients with cerebral palsy when the arm is not fully extended due to a smaller lever arm at the shoulder, which makes sense because forward trunk flexion decreases the antigravitational torque at the shoulder in the final posture ^13^. Fuller *et al.* (2013) suggest a similar hypothesis for healthy subjects who begin to use trunk flexion after a fatigue protocol: Using the trunk would relieve the shoulder muscles by recruiting the trunk muscles. Hence, it is likely that the trunk is recruited to limit the torque at the shoulder joint in the final posture. Doing so, the objective of the task would be pursued (*i.e.* bringing the hand closer to the target), while the effort would be better distributed and with greater torque reserve for the involved joints.

In this paper, we hypothesise that trunk recruitment becomes significant when the shoulder is too close to maximal voluntary torque. We test this hypothesis by comparing the final posture when the hand is unloaded versus when the hand is loaded at 75% of the maximum antigravity shoulder torque.

## 2. Methods

### 2.1. Participants

Twenty-six healthy participants (12 males, age 21 ± 3 years, 3 left-handed, height 1.73 ± 0.09 m, weight 66.92 ± 9.29 kg, maximum voluntary force (MVF) of shoulder flexion 65.32 ± 19.50 N.m) participated in this study. Participants were excluded if they had shoulder pain or any other problem that could affect the reaching task. Written informed consent was obtained from all participants before their inclusion. This study was performed in accordance with the 1964 Declaration of Helsinki. The Institutional Review Board of EuroMov at the university of Montpellier approved the study (IRB-EM 1901C).

### 2.2. Procedure

Participants had to reach a target with the side of their thumb nail. The target was table tennis ball fixed in front of the participant at a height of 80 cm ^14^. The distance to the target was adjusted to correspond to the length of the participant’s active stretched arm measured from the medial axila to the distal wrist crease ^15^. The height of the chair was adjusted so that the target horizontal projection was between the participant’s navel and xiphoid process.

In the starting position, the participants were seated with their feet on the floor, their backs in contact with the back of the chair and their forearms on the armrest. The reaching movement was assessed in 2 × 2 conditions: spontaneous trunk use or restrained trunk use × hand unloaded or hand loaded. In the spontaneous trunk use condition, participants had to reach the target at a natural pace, wait 1 second and return to the starting position. In the restrained trunk use condition, participants had to reach the target while minimising trunk movement: the experimenter manually applied a light proprioceptive feedback on the participant’s shoulders, as a reminder to minimise trunk movement. We did not use a belt to restrain the trunk in order to leave the participant free to use the trunk if necessary, and thus avoid task failure.

In both trunk conditions, participants performed 5 unloaded trials and 5 loaded trials, i.e., carrying a dumbbell (figure 1). The assessed hand was chosen pseudo-randomly (12 left, 14 right) so that half of the participants performed the task with their dominant hand, and the other half with their non-dominant hand. In the hand unloaded condition, the weight of the arm without the dumbbell corresponded to 13.0% (±3.7%) of the MVF of the shoulder. In the hand loaded condition, the weight of the dumbbell was set as a function of the MVF of the participant, in order that the weight of the arm including the dumbbell corresponded to 75.0% (±5.5%) of the MVF of the shoulder.

**Figure 1.**
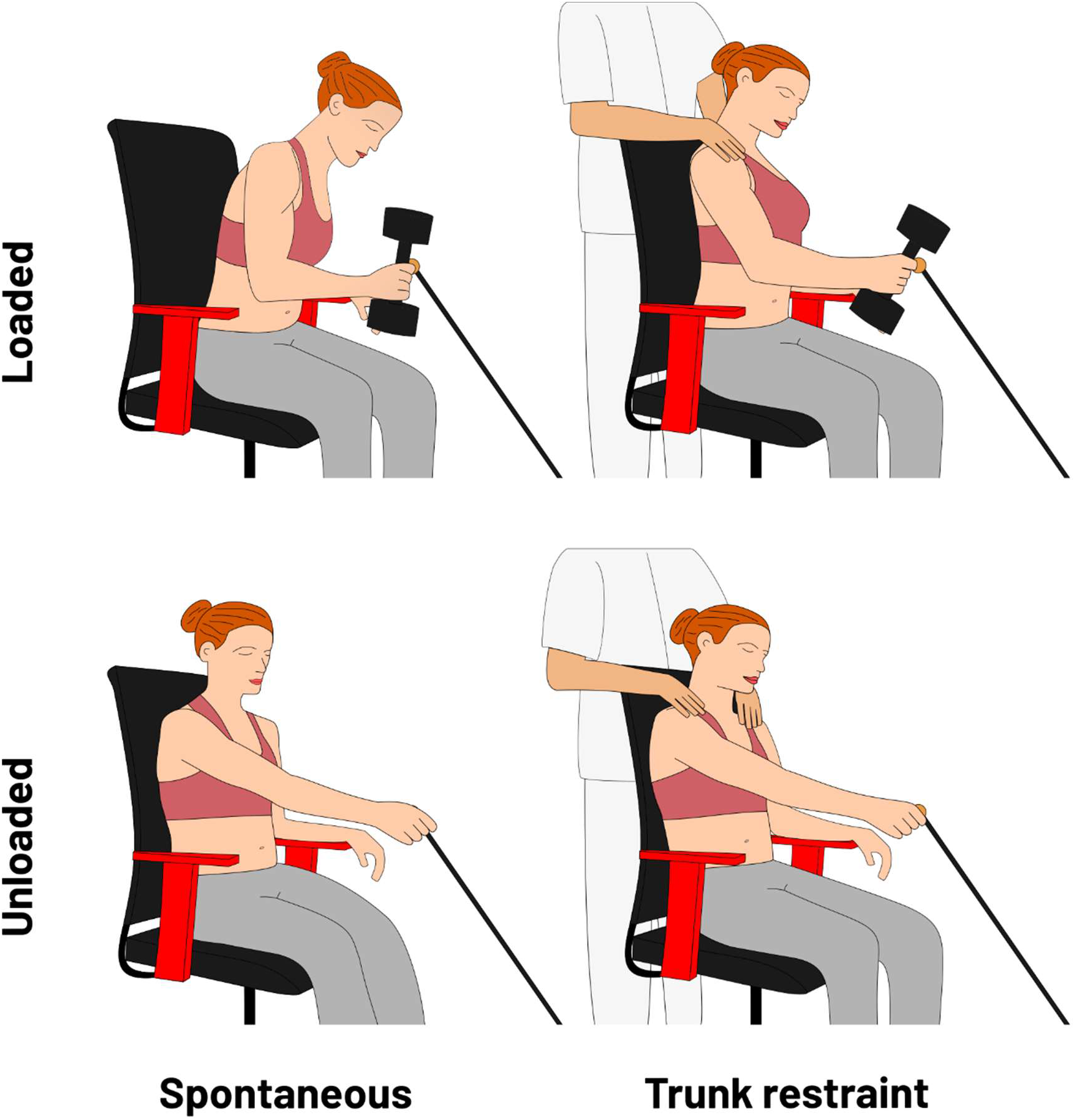
Schema of the four experimental conditions. In the loaded conditions (upper row), participants held a dumbbell which mass corresponded to 75% of their maximal voluntary shoulder force (MVF). In the unloaded conditions (lower row), participants had to reach the target without the dumbbell. In the spontaneous conditions (left column), participants had to reach the target in a natural way. In the trunk restraint conditions (right column), participants had to reach the target while minimising trunk movement.

The MVF of the shoulder was the maximum torque producible by the shoulder during an isometric shoulder flexion task against gravity, including arm weight. Seated participants had to hold a 2 kg dumbbell in front of them with the arm extended and pull up as much as possible for 3 seconds. The dumbbell was linked with a static rope to a force sensor attached to the ground. Participants had to stay in contact with the backrest during the entire contraction. Any help of the contralateral arm was prohibited. Experimenter encouraged the participant verbally. We retained the maximum score over 3 trials separated with a rest time of 1 minute. Maximum antigravity shoulder torque was calculated based on the recorded maximum force and lever arm, plus the torque due to the weight of the arm during the task.

### 2.3. Experimental setup

Movements were recorded at 100 Hz with 8 infrared optical cameras from the Vicon Motion Capture System (Vantage V5, lens 8.5 mm, Oxford Metrics, UK). VICON Nexus 2 software was used to save time series of each marker. The experimenter placed markers on the target and the manubrium, and for each body side on the 1^st^ metacarpal, the lateral epicondyle of the humerus, the acromion process, and the iliospinal. For each side, we corrected iliospinal marker position before data analysis to correspond at best to the anatomical centre of hips joints. Markers of both sides were needed to compute shoulder and trunk angles.

### 2.4. Kinematic analysis

Data analysis was performed with SciLab 6.0.2.

First, all position time series were low pass filtered at 5 Hz with a dual pass second order Butterworth filter.

Second, we computed the beginning and end of the reach. Because the goal of a reaching task is to bring the hand to the target, that is, to reduce the hand-to-target distance, what is important for task success is the hand-to-target Euclidean distance. The hand-to-target Euclidean distance summarises the 3D effector space into a 1D task space (where movement matters) leaving aside a 2D null space (where movement does not impact task success). We fixed the beginning of the movement (t_0_) when the Euclidean velocity of the hand in task space became positive and remained positive until the maximum velocity. The end of the movement (t_final_) was when the Euclidean distance to the target reached its minimum.

Third, we computed the trunk recruitment as a percentage of reach length. Reach length is the hand-to-target Euclidean distance at t_0_. In order to know the normalised implication of the trunk to the reach, we computed trunk recruitment as the ratio between shoulder displacement and reach length. More precisely, trunk recruitment = (shoulder to target distance at t_0_ − shoulder to target distance at t_final_) / (hand to target distance at t_0_ − hand to target distance at t_final_). Over the 5 trials, the trial with the median trunk recruitment was retained for data analysis.

Fourth, antigravity shoulder torque was computed as the Euclidean norm of the 3D static torque against gravity at the final posture. In order to compute the static torque, the positions of centre of mass and the absolute weight of upper limbs were approximated from the height and weight of participants following De Leva’s equations (1996). Our model considers the torques applied to the shoulder by the upper arm (shoulder to elbow link), the forearm and hand (elbow to hand link) and the dumbbell, as described in the equation 1.

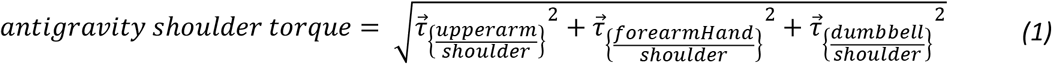

The torque applied to the shoulder by each component of the model was calculated as the cross product of the lever arm vector and the gravity vector of the component (equation 2).

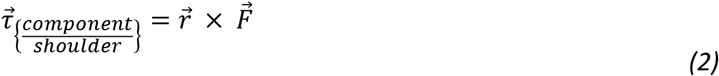

Where 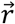 is the lever arm vector (a position vector from the shoulder to the centre of mass of the component) and 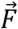 is the gravity vector of the component. The equation can be developed as:

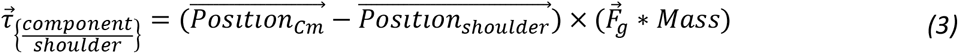

Where 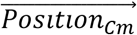 is the 3D position of the centre of mass of the component, 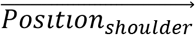 is the 3D position of the shoulder joint, 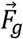 is the force of gravity and *Mass* is the mass of the component.

### 2.5. Statistical analysis

The statistical analysis was conducted using R 3.6.1 with rstatix and stats packages. Density plots and quantiles/quantiles plots revealed a violation of the normality assumption inside data groups. Therefore, all hypotheses were tested in a non-parametric way, using paired samples Wilcoxon signed rank tests. Because of the multiple comparisons, p-values were corrected by the method of Benjamini & Hochberg (1995) with a false discovery rate (FDR) of .05. This method aims to control the expected proportion of false discoveries amongst the rejected hypotheses (the FDR) in order to reduce type I errors while limiting type II errors. The FDR method is less stringent than the family-wise error rate and is therefore more powerful than a simple Bonferroni-type procedure. The significance level was set at .05 in all analyses. The effect size r was calculated as the Z statistic divided by the square root of the number of observations ^18^. Data are presented as median (± interquartile range).

## 3. Results

### 3.1. Increasing the weight of the arm raises trunk recruitment

Reaching in the spontaneous unloaded condition induced a trunk recruitment of 11% of reach length. Trunk recruitment increased to 27.5% when loaded (W = 0, p < .001, r = .87). In the trunk restraint conditions, trunk recruitment could be reduced to 8% when unloaded (W = 22, p < .001, r = .72), and to 12% when loaded (W = 0, p < .001, r = .87) (figure 2a). The increase in trunk recruitment was not different between subjects with the dominant side loaded and subjects with the non-dominant side loaded (W = 97, p = .54, r = .13). These results indicate that handling a heavy dumbbell during reaching increases trunk recruitment (table 1). When asked, all participants were able to reduce trunk recruitment in the loaded condition.

**Table 1.** Differences in angular rotation and raw displacements between final and initial posture. When the hand is unloaded and the trunk is restrained, the coordination is mainly based on elbow and shoulder use (left of the table). When the hand is loaded or the trunk is free, trunk recruitment increases, which reduces elbow and shoulder use (right of the table).

| Trunk<br>Hand | Trunk restraint<br>Unloaded | Spontaneous<br>Unloaded | Trunk restraint<br>Loaded | Spontaneous<br>Loaded |
| --- | --- | --- | --- | --- |
| Elbow extension (°) | 43 (±19) | 37 (±13) | 29 (±16) | 14 (±19) |
| Shoulder flexion (°) | 42 (±10) | 41 (±12) | 39 (±12) | 31 (±15) |
| Shoulder adduction (°) | 22 (±11) | 19 (±8) | 11 (±12) | 12 (±8) |
| Trunk flexion (°) | 0 (±0) | 1 (±0) | 1 (±2) | 5 (±5) |
| Trunk rotation (°) | 4 (±2) | 6 (±3) | 12 (±6) | 17 (±6) |
| Hand displacement (mm) | 314 (±46) | 314 (±42) | 317 (±51) | 338 (±62) |
| Shoulder displacement (mm) | 25 (±13) | 36 (±16) | 63 (±40) | 138 (±86) |
| Trunk displacement (mm) | 5 (±4) | 11 (±5) | 38 (±25) | 79 (±73) |

**Figure 2.**
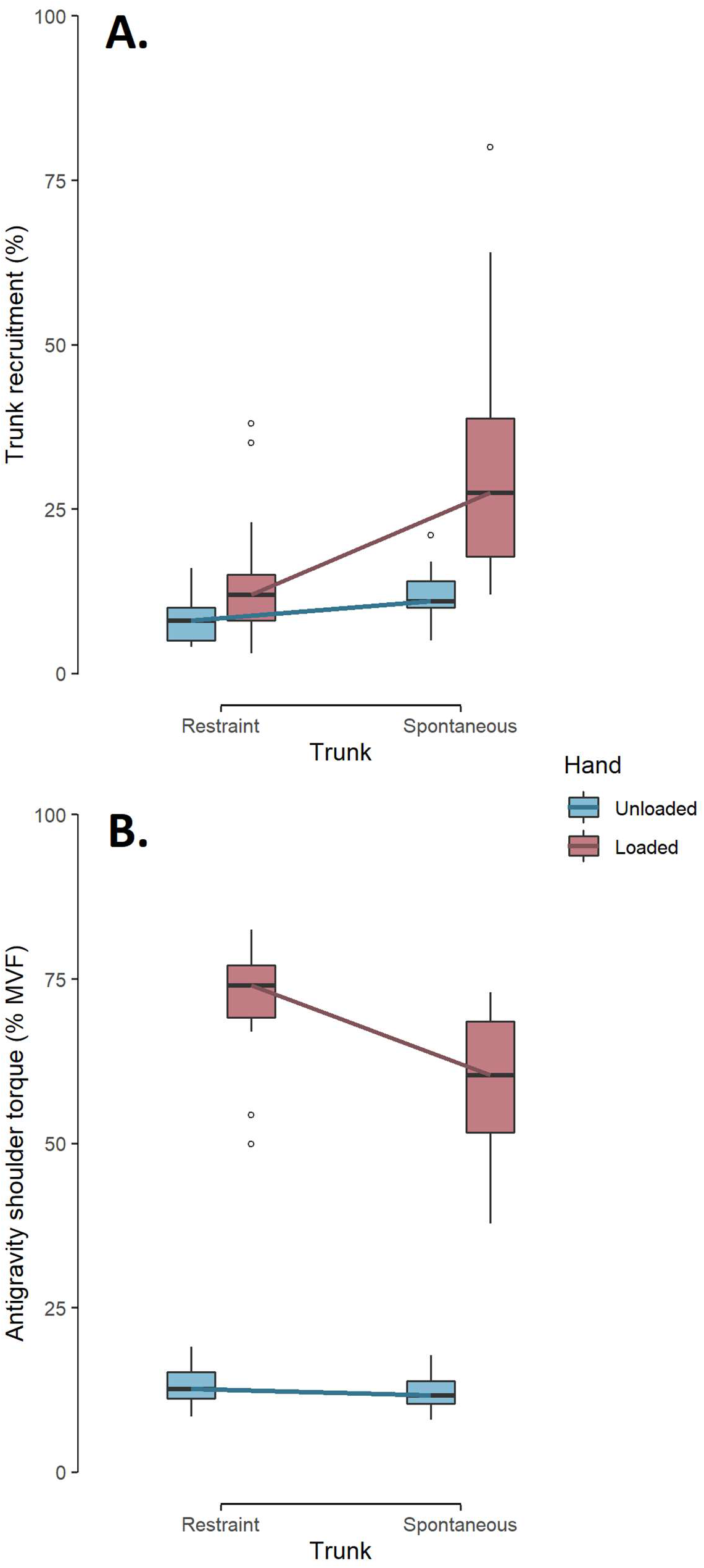
Trunk recruitment (Panel A) and antigravity shoulder torque (Panel B) as a function of trunk restraint and hand load. Trunk recruitment is in percentage of reach length. Antigravity shoulder torque at the final posture is in percentage of maximal voluntary shoulder torque. In each boxplot, the internal horizontal bar represents the median, boxes extend from first to third quartile, whiskers extend from minimal to maximal values excluding outliers and white circles represent outliers. All comparisons are statistically significant. The figure shows that trunk recruitment stays low in the unloaded conditions. When the hand is loaded, trunk recruitment is doubled, and antigravity shoulder torque is decreased by 18% in the spontaneous condition.

### 3.2. Trunk recruitment reduces antigravity shoulder torque

Trunk restraint increased shoulder antigravity torque in the final posture from 11.7% to 12.7% of maximum when the hand is unloaded (W = 351, p < .001, r = .87) and from 60.4% to 74.1% of maximum in the loaded condition (W = 349, p < .001, r = .86) (figure 2b). The decrease in shoulder torque was not different between subjects with the dominant side loaded and subjects with the non-dominant side loaded (W = 92, p = .72, r = .08). In other words, trunk restraint increased the mechanical cost that is needed at the shoulder to maintain the final posture. Given the increased arm weight in the loaded condition, we made no comparison between unloaded and loaded conditions.

## 4. Discussion

We tested the hypothesis that, when reaching to grasp an object from a seated position, people recruit the trunk when shoulder torque is too close to maximal. We found that when the hand is loaded to 75% of the maximum antigravity shoulder torque, trunk recruitment increases from 11.0% to 27.5%. By flexing and rotating the trunk, participants spontaneously reduced antigravity shoulder torque from 74.1% to 60.4% of maximum voluntary torque.

### 4.1. Trunk is a crucial component of the reaching coordination

The spontaneous unloaded condition informs about the final posture that is naturally selected when reaching to grasp an object from a seated position. From a side view, the task requires to move the hand forward. To do this, people combine elbow extension (43°) and shoulder flexion (42°), with no flexion of the trunk (0°). From a frontal view, the task requires moving the hand towards the midline of the body. To do this, people combine shoulder adduction (22°), with a small trunk rotation (4°) (see Table 1 and Figure 3, top row)).

**Figure 3.**
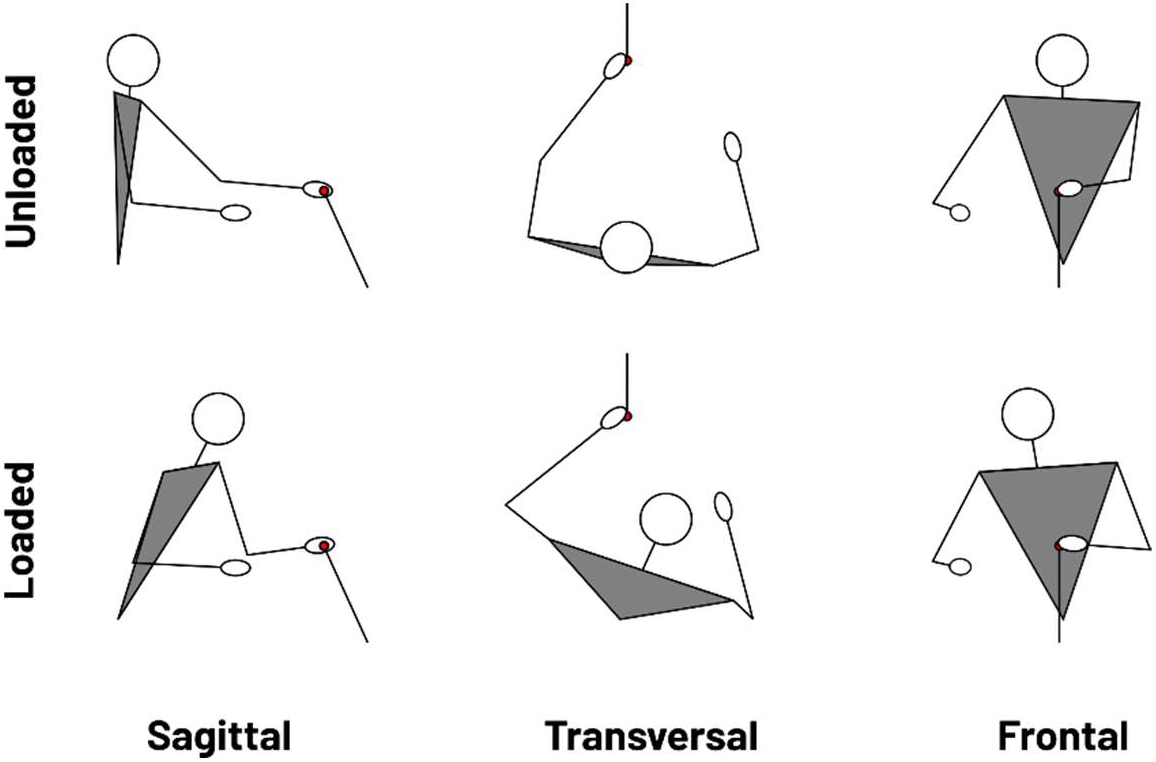
Final posture of a typical participant in the spontaneous condition as a function of load (Unloaded: top row, Loaded: bottom row). Diagrams are viewed from the side (sagittal plane, left column), viewed from above (transverse plane, middle column) and viewed from the front (frontal plane, right column). The red circle represents the target. When the hand is loaded, the participant has less elbow extension, less shoulder flexion and shoulder abduction at the expense of greater trunk flexion and rotation. The figure shows that the greater recruitment of the trunk brings the shoulder closer to the target, resulting in less gravitational torque at the shoulder, as can be inferred by comparing the sagittal views.

In the spontaneous unloaded condition, participants use a combination of elbow extension and shoulder flexion and adduction, but hardly recruit the trunk (figure 3, top row). This “arm strategy” is in agreement with the theory of optimal control, which says that human movement minimise costs ^3^. Indeed, although participants could recruit the trunk, they do not because this would imply moving a large and heavy segment, and therefore to mobilise large muscles. It is more economical to mainly use the arm and forearm segments, which are less heavy and have less inertia than the trunk.

In the spontaneous loaded condition, participants still use a combination of elbow extension and shoulder flexion and adduction, but they do recruit the trunk (figure 3, bottom row). This “trunk strategy” decreases shoulder-elbow participation, and therefore minimises antigravity shoulder torque (Figure 2). Indeed, when the arm is loaded and the trunk restricted, antigravity shoulder torque of the final posture is 74% of the maximum, which is a high intensity. According to Diedrichsen *et al.* (2010), when muscles are strongly activated, metabolic processes are not optimised and therefore the need for adenosine triphosphate (ATP) to maintain exertion is high. In addition, signal-dependent noise increases with the intensity of activation ^19^, which may affect the precision of the reach. Thus, when the arm is loaded, the “arm strategy” likely involves a more energy-consuming and imprecise movement. Participants therefore change their coordination and their final posture so to reduce antigravity shoulder torque. The “trunk strategy” distributes the work over more effectors and thus reduces the load on shoulder muscles. The “trunk strategy” improves metabolic processes and reduces signal-dependent noise when the weight of the arm is high relative to the strength of the shoulder. This gain probably outweighs the metabolic expense required to recruit trunk muscles.

Diedrichsen et al. (2010) rename this principle “distribution of work across multiple effectors”. The authors consider that the cost of the task is equal to the squared sum of the motor commands. Indeed, this cost function is well correlated with ATP consumption and therefore with the effort required. According to this cost function, by penalising high activations, the distribution principle would minimise the total cost of the movement, and therefore minimise the required effort. Our results are consistent with the conclusions of Wang & Dounskaia (2012). They show that increasing arm weight accentuates, and thus makes visible the minimisation of muscular effort. Thus, when the arm is unloaded, effort minimisation is still at work but is less noticeable.

Here, we confirm the idea of Peeters *et al.* (2018) and Fuller *et al.* (2013) that trunk movement occurs when the shoulder has to develop a too great strength in relation to its capacity. This means that in addition to stabilising the arm and regulating the reachable distance ^8,21^, trunk movements emerge to avoid high shoulder torques.

#### Limitation

Participants did the reaching movement without holding a dumbbell in the unloaded condition. So, it is possible that some of the coordination differences are due to the difference in hand orientation between the dumbbell (loaded) and the non-dumbbell (unloaded) condition. However, in order to avoid this bias, the target was designed not to interfere with the movement performed with the dumbbell, and participants were instructed to hit the target with the side of their thumb, which theoretically produces a strictly similar movement, whether or not the participant is holding the dumbbell.

### 4.2. Implications for motor control

The theory of optimal control states that movement coordination is performed to meet the constraints of the task while minimising a cost function ^22^. Objective cost functions were first explored to minimise a mechanical cost ^23,24^, but according to Loeb (2012), cost functions not based on subjective feelings make no sense in a neurobiological system. More human-oriented functions were then developed, such as minimum neuromuscular effort ^26^, minimum metabolic cost ^27,28^ or maximum comfort ^29^.

Finally, other functions combine objective and subjective effort ^30^ or introduce modulation by individual traits ^4,31^. In the face of this diversity, there is currently no consensus on the list of ingredients to include in the cost function.

Here we show that in a seated reaching task, when the arm is heavy relative to the shoulder strength, healthy participants modify their final posture to avoid excessive antigravity torque at the shoulder. By recruiting the trunk in the loaded condition, the antigravity torque at the shoulder decreases from 40% to 25% of maximum. Although this result is consistent with the predictions of the optimal control framework, future work is necessary to ensure that the observed strategy is optimal.

Instead of performing optimally, biological organisms may just do “good enough” ^25^. Indeed, muscle coordination might be habitual rather than optimal in a wide variety of situations, which would explain the absence of consensus on the nature of the cost function. Following this approach, a disturbed sensorimotor system will more probably exhibit, at first, the habitual coordination adapted to the new constraints instead of the optimal solution that might come with learning ^32^. In the present experiment, when loaded, participants immediately performed significant trunk recruitment. This suggests that participants have already learned that the habitual coordination is no longer “good enough” when the torque at the shoulder becomes excessive.

The strategy of coordination is directly adapted to the constraints to avoid a shoulder torque greater than about 60%, i.e. to guarantee a shoulder torque reserve of about 40%. When pushed to their maximum in the trunk restraint condition, people voluntarily reduce the reserve to about 25% (figure 4, pink bars). In the trunk restraint condition, one would have expected that trunk recruitment would tend towards zero. Yet, participants chose to retain some trunk recruitment, resulting in 25% unmobilised shoulder reserve. This suggests that the spontaneous 40% reserve might be the sum of a mobilisable comfort reserve of 15%, and a non-mobilisable safety reserve of 25%. Keeping a reserve of strength is tantamount to the safety margin when grasping an object ^33^, and more generally, keeping a safety reserve is a behavioural strategy adopted by humans to be able to react to unexpected perturbations, which is mandatory to succeed in the task every time.

**Figure 4.**
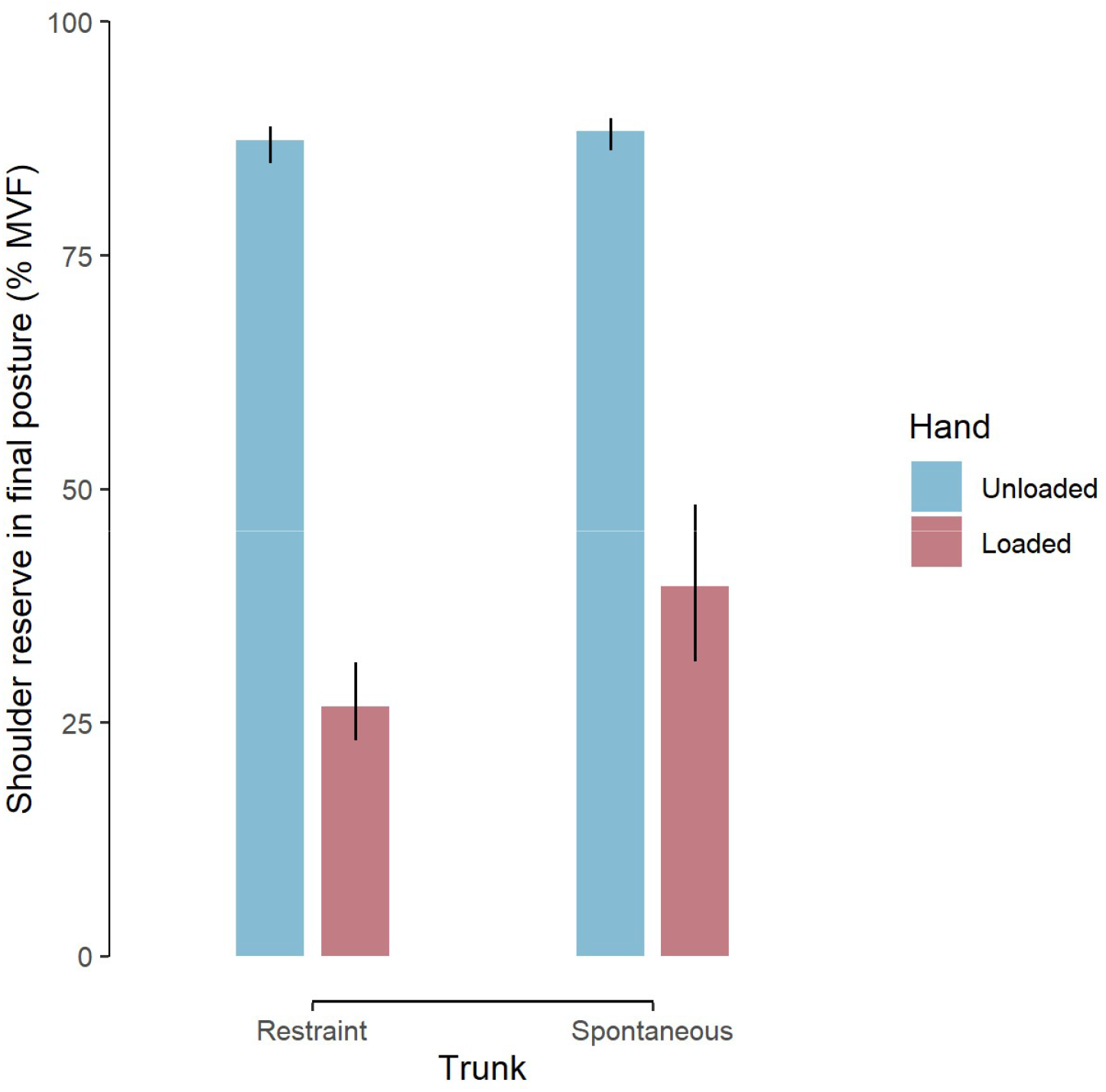
Shoulder reserve in final posture as a function of trunk restraint and hand load. Shoulder reserve is in percentage of maximal antigravity shoulder torque. In each barplot, the top of the bar represents the median, error bar extends from the first to the third quartile. All comparisons are statistically significant. The figure shows that shoulder reserve stays high in the unloaded conditions. When the hand is loaded and the trunk is restrained, shoulder reserve is considerably reduced.

Although most of the current cost functions are linked to human effort, they are generally derived from experiments where participants spontaneously perform far from their limits ^31,34^. Here we demonstrate that, to better model the reaching movement where people are pushed towards their maximum, realistic cost functions should penalise too strong activations. Our results suggest that a reserve of about 40% of maximal force is the limit of acceptable activation without spontaneously recruiting a novel degree of freedom.

#### Limitations and future work

If this work lays the foundations for understanding the activation threshold beyond which the effort is avoided, it remains preliminary. First, the antigravity torque at the shoulder is only estimated statically for the final posture, which does not fully reflect the actual activation of the shoulder muscles. Torques could be estimated dynamically during the reach and muscle activations could be investigated using EMG analysis. However, with a motion time of 1.97s (±0.6) and a dwell time of at least 1s at the target, the antigravitational torque at the final posture is the main constraint (e.g., an order of magnitude greater than the maximum dynamic torque ^35^). This results in greater activation of the anterior deltoid at the end of movement in slow reaches ^36^. Second, further experiments could measure how trunk recruitment increases as a function of shoulder torque, for example to get a picture of the non-linearity of this relation. Third, the reaching task studied here is very stereotypical. Further work could measure the evolution of coordination in more ecological tasks, with greater or lesser spatial accuracy requirements.

### 4.3. Perspectives for stroke research

Our result that trunk recruitment is a cost-effective solution when shoulder torque is too close to maximal has consequences for the interpretation of movement disorders. The most disabling factor after a stroke is muscle weakness in the hemiparetic arm ^37^. In other words, after a stroke, the weight of the arm is too heavy for the available strength of the shoulder muscles ^38,39^. As a result, people with stroke are close to maximum neuromuscular activation during a reaching movement ^40^. After a stroke, it is likely that the principles governing sensorimotor control and learning are the same ^41,42^. Hence, with the very same control policy in the face of an atypical body, the emerging coordination is atypical ^43^. By recruiting the trunk, patients follow the typical control policy: they reduce the necessary shoulder torque and thus minimise effort. This interpretation is consistent with empirical results showing that arm weight support decreased neuromuscular activation of the shoulder ^44,45^, and so increased the reachable space of the patients ^46,47^. This confirms that the ability to lift the arm is a strong limiting factor when reaching after a stroke.

When the available force is limited, it may be more efficient to freely use all the available degrees of freedom instead of spending more attention, effort, energy to perform the task more “normally” (*i.e.* with the “arm strategy”). Thus, trunk compensation by stroke victims is not ineluctably a “bad synergy” resulting from a pathological synergy or from an impaired control ^48,49^, but could instead testify that stroke movement continue to follow optimal control principles, but with different constraints (such as reduced maximum shoulder torque or greater noise), as claimed in other diseases ^43^. Further research is needed to confirm that both people with and without stroke recruit their trunk when the arm is too heavy for the available shoulder force. For example, experimentally reducing the weight of the arm of post-stroke individuals should result in a proportional reduction in trunk compensation. If so, increasing antigravity shoulder force might be a way to facilitate arm use in individuals with stroke. Thus, it may be worth considering training that emphasises strength in addition to conventional therapies ^50^.

However, care must be taken not to confuse the strength reserve with what we call a “coordination reserve”. The latter is a form of neuroplastic cognitive reserve that explains “the disjunction between the degree of brain damage and its clinical outcome” ^51^ and that can be drawn upon to change the coordination through learning ^52,53^.

## 5. Conclusion

Our experiment shows that, in young healthy persons reaching at a target while seated, trunk recruitment becomes visible when the participant faces antigravity force limitations at the shoulder. The recruitment of the trunk is a cost-effective solution to keep the activation of the shoulder muscles under 60% of their maximum. More generally, our results suggest that all joints in the kinematic chain actively participate in the movement, but that the amplitude of their recruitment is mediated by force constraints: energy-consuming joints are recruited only when necessary, to minimise the use of the force reserve of the other joints.

## Data availability

The raw datasets generated during the current study are available in the Open Science Framework repository: https://osf.io/r3xcu/

## Code availability

The code generated during the current study is available in the Open Science Framework repository: https://osf.io/r3xcu/

## Acknowledgements

This study was supported by the LabEx NUMEV (ANR-10-LABX-0020) within the I-SITE MUSE. We thank Victor Barradas and Nicolas Schweighofer for their comments on the article. We acknowledge Germain Faity for drawing the schema of the experimental conditions.

## Authors’ contributions

GF designed the protocol, recorded and analysed the data, wrote the first version of the manuscript. DM and JF assisted GF in designing the protocol, analysing the data and provided guidance for improving the manuscript. SP assisted GF in setting up the protocol, recording the data and improving the manuscript. All authors accepted the latest version of the manuscript.

## Additional information

### Competing interests

The authors declare that they have no competing interests.

## Notes

https://osf.io/r3xcu/

### Competing Interest Statement

The authors have declared no competing interest.

### Summary of Updates

In this version, we have toned down the conclusions made regarding optimal control and we have adapted the manuscript accordingly. We have also mitigated the perspectives for stroke research by removing the consequences for therapists from the manuscript. Finally, we added a figure (Fig 3) to better illustrate the changes in final posture as a function of arm weight.

